# Characterizing cellular heterogeneity in chromatin state with scCUT&Tag-pro

**DOI:** 10.1101/2021.09.13.460120

**Authors:** Bingjie Zhang, Avi Srivastava, Eleni Mimitou, Tim Stuart, Ivan Raimondi, Yuhan Hao, Peter Smibert, Rahul Satija

**Author notes:** These authors contributed equally (Bingjie Zhang, Avi Srivastava).

## Abstract

New technologies that profile chromatin modifications at single-cell resolution offer enormous promise for functional genomic characterization. However, the sparsity of these measurements and the challenge of integrating multiple binding maps represent significant challenges. Here we introduce scCUT&Tag-pro, a multimodal assay for profiling protein-DNA interactions coupled with the abundance of surface proteins in single cells. In addition, we introduce scChromHMM, which integrates data from multiple experiments to infer and annotate chromatin states based on combinatorial histone modification patterns. We apply these tools to perform an integrated analysis across nine different molecular modalities in circulating human immune cells. We demonstrate how these two approaches can characterize dynamic changes in the function of individual genomic elements across both discrete cell states and continuous developmental trajectories, nominate associated motifs and regulators that establish chromatin states, and identify extensive and cell type-specific regulatory priming. Finally, we demonstrate how our integrated reference can serve as a scaffold to map and improve the interpretation of additional scCUT&Tag datasets.

## Introduction

Technologies that enable unsupervised transcriptomic profiling at single-cell resolution (i.e. scRNA-seq) represent powerful tools not only for the discovery of cell types and states, but also to reveal the structure and function of transcriptional regulatory networks^1–6^. In the same way, new methods for single-cell chromatin profiling, such as scATAC-seq, can lead to the identification and characterization of individual genomic regulatory regions, and to explore their variation in a heterogeneous population^7,8^. Assays that profile chromatin accessibility exhibit a largely binarized phenotype - partitioning genomic regions into accessible or inaccessible elements in different tissue and cellular contexts. As a complement to accessibility measurements, genome-wide histone modification profiles offer an exciting opportunity to segment the genome into a more nuanced set of functional elements^9–12^. The “histone code hypothesis” proposes that the combinatorial presence of multiple histone modifications marking a genomic region serves as a proxy for its regulatory function^13,14^.

Techniques like ChIP-seq and CUT&Tag have been widely applied to measure the binding profiles of multiple transcription factors and histone modifications^15,16^. In addition, tailored computational tools such as ChromHMM^17^ and Segway^10^ can integrate the combinatorial binding patterns defined by each mark, and assign individual genomic elements to a set of learned functional ‘states’. Recently, multiple studies have demonstrated the ability to measure CUT&Tag profiles in single cells^16,18–21^. This represents an opportunity to not only identify regions that exhibit cell type-specific accessibility, but also to highlight elements whose acquisition of activating, repressive, or heterochromatic signatures varies within a heterogeneous population.

These advances lay out an exciting challenge for single cell genomics: can the binding profiles of multiple histone modifications be used to infer the function of any individual genomic element, within a single cell? While the availability of scCUT&Tag takes an important step towards this vision, two key challenges remain. First, single-cell chromatin data remains extremely sparse, particularly due to the inherent challenges associated with limited DNA input molecules. Second, scCUT&Tag experiments typically measure only a single histone modification profile in a single cell. While informative, only the integrated combination of multiple histone modifications can be used to annotate functional state, but no current technology can robustly and simultaneously profile many histone modifications in the same cell, particularly when these marks may exhibit overlapping binding patterns. Recent pioneering advances enable paired measurements of cellular transcriptomes and individual CUT&Tag profiles using custom combinatorial-indexing based workflows^22,23^, and demonstrate how multimodal technologies can facilitate integrative analysis.

Here, we present two new techniques aimed to address these key challenges. We first introduce scCUT&Tag-pro, a multimodal single-cell technology, that enables simultaneous profiling of individual CUT&Tag profiles paired with surface protein abundances at single-cell resolution. Our approach is compatible with the widely used 10x Genomics Chromium system and complements recently introduced technologies for simultaneous CUT&Tag and transcriptomic profiling that leverage custom combinatorial indexing workflows^22,23^. Second, we introduce a downstream analysis strategy for multimodal datasets: scChromHMM. Our method first integrates data from multiple scCUT&Tag-pro experiments together into a common manifold, computationally generating co-assay profiles for six histone modifications within individual cells. These measurements are used as input to a single-cell extension of the ChromHMM algorithm, in order to return chromatin state annotations for each 200 base pair (bp) genomic window at single-cell resolution.

We apply our tools and technologies to generate 64,876 scCUT&Tag-pro profiles (Supplementary Table 1) and combine this with existing datasets to create an integrated multimodal atlas of human peripheral blood mononuclear cells (PBMCs) that encompasses nine modalities spanning the central dogma. Together, these technologies and analytical tools allow us to explore heterogeneity in genome function at single-cell resolution, identify sequence elements whose presence accompanies changes in chromatin state, and project new datasets onto our multimodal reference. We propose that these tools will help to facilitate the analysis and interpretation of chromatin state heterogeneity in single cells, and lead to a better understanding of its role in establishing and regulating cellular identity.

## Results

### Simultaneous profiling of surface proteins and CUT&Tag profiles in single cells

The combinatorial pattern of post-translational modifications present on histone proteins correlates with the chromatin structure and regulatory potential of a genomic locus^13,14,24,25^. We reasoned that multimodal single-cell technologies could help address challenges in scCUT&Tag analysis by coupling robust single-cell measurements of one modality with more sparse measurements from another. For example, CITE-seq/REAP-seq^26,27^, and ASAP-seq/ICICLE-seq^28,29^ pair simultaneous measurements of highly expressed and well-characterized cell surface proteins with unsupervised but sparser transcriptomic measurements or chromatin accessibility profiles. We and others have recently demonstrated that the protein information in CITE-seq is highly informative in determining cellular state^30–32^, while the paired RNA-seq or ATAC-seq measurements enable the characterization of gene regulatory networks. Importantly, ASAP-seq and ICICLE-seq are enabled by recent optimizations in cell fixation and permeabilization that enable ATAC-seq to be performed on whole cells instead of individual nuclei^33^.

We propose that a similar multimodal strategy could be applied for genome-wide profiling of protein-DNA interactions as well. We developed scCUT&Tag-pro, an assay that performs CUT&Tag assays^16^ on whole cells while simultaneously measuring surface protein levels (Figure 1A; Supplementary Methods). Inspired by ASAP-seq^28^, our fixation and permeabilization conditions retain the cell membrane and associated proteins, enabling us to simultaneously measure cellular immunophenotypes (Figure 1A). Briefly, cells are first stained with oligo-conjugated antibodies. Subsequently, monovalent Fab fragment is added in order to block the potential binding of proteinAG to these antibodies. Cells are then lightly fixed with 0.1% formaldehyde and permeabilized with an isotonic lysis buffer. We found that removing Digitonin from all buffers substantially alleviated cell ‘clumping’ issues which have been previously described^18^. We utilize an antibody against a histone modification of interest to direct the proteinAG-Tn5 fusion protein to marked genomic sites and initiate site-specific tagmentation using a high-salt buffer containing magnesium. Tagmented cells can then be used as input for library preparation using the 10x Genomics scATAC-seq kit, followed by sequencing of both CUT&Tag as well as antibody-derived tag (ADT) libraries. We note that scCUT&Tag-pro is also compatible with antibody-mediated Cell Hashing^34^, allowing us to increase the throughput of each experiment.

**Figure 1:**
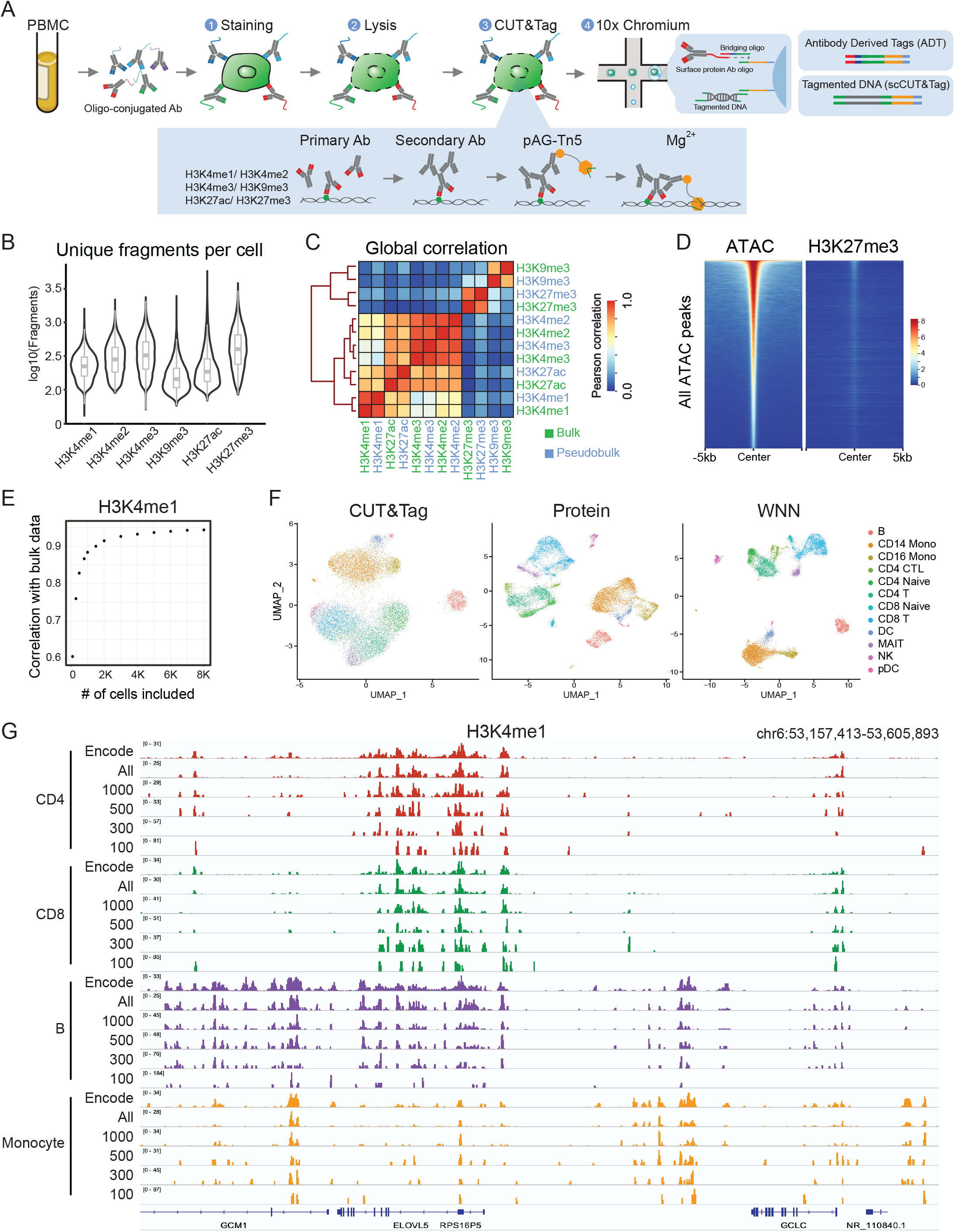
scCUT&Tag-pro enables simultaneous profiling of CUT&Tag and protein levels. (A) Schematic of experimental workflow, which is compatible with the 10x Chromium system. **(B)** Distribution of unique fragments obtained per-cell for six histone modifications, profiled in separate PBMC scCUT&Tag-pro experiments. **(C)** Pseudobulk profiles of scCUT&Tag are well-correlated with bulk CUT&Tag profiles of human PBMC. **(D)** Tornado plots of genomic regions ordered by chromatin accessibility. We observe no enrichment of H3K27me3 in accessible regions, indicating minimal open chromatin bias. **(E)** Relationship between the number of cells included in a pseudobulk CUT&Tag profile of human PBMC, and the pearson correlation with a bulk experiment. **(F)** UMAP visualizations of 12,770 single cells profiled with H3K4me1 scCUT&Tag-pro and clustered on the basis of CUT&Tag profiles, cell surface protein levels, and weighted nearest neighbor (WNN) analysis which combines both modalities. Cluster labels are derived from WNN analysis. **(G)** Comparing pseudobulk CUT&Tag-pro profiles with ChIP-seq data from ENCODE. Including all cells assigned to each cell type results in pseudobulk tracks that closely mirror ENCODE profiles. However, even when downsampling to 300 cells per cluster, cell type-specific patterns can still be observed.

In this study, we focused on acquiring scCUT&Tag-pro data from healthy human PBMC and aimed to study the genome-wide localization patterns for six histone modifications (H3K4me1, H3K4me2, H3K4me3, H3K27ac, H3K27me3, and H3K9me3). In each experiment (Supplementary Table 1), we profiled a single histone modification, alongside an optimized panel of 173 antibodies from the Biolegend TotalSeq-A catalog (Supplementary Table 2). In total, we collected scCUT&Tag-pro from 64,876 individual cells.

We first evaluated the sensitivity and specificity of our CUT&Tag profiles. For example, in our H3K27me3 dataset we observed, on average, 501 unique fragments per cell with an 84.1% uniquely mapping rate (Supplementary Figure 1A), comparable to but slightly below a recent pioneering study^19^ introducing scCUT&Tag on single nuclei from the same system (802 average unique fragments/cell, Figure 1B, Supplementary Figure 1B). Importantly, for all marks, we observed that a pseudobulk profile generated from our single-cell data closely mirrored simultaneously generated bulk CUT&Tag profiles from PBMC (Figure 1C). Moreover, we found that regions marked by H3K27me3 exhibited minimal overlap with accessible regions identified in a scATAC-seq dataset^28^, demonstrating a lack of ‘open-chromatin bias’ that can be present when CUT&Tag is performed in low-salt conditions^16^ (Figure 1D). Through downsampling analysis, we found that the quantitative accuracy of each histone modification pseudobulk profile was dependent on the number of included single cells but began to saturate between 500-1,000 cells (Figure 1E). Moreover, we found that unsupervised clustering of each scCUT&Tag dataset using an LSI-based dimensional reduction workflow (Supplementary Methods) was capable of identifying major cell groups that define the human immune system (Figure 1F), but did not recapitulate the high-resolution identification of cell states as observed in scRNA-seq or CITE-seq data^30^.

### Cell surface protein measurements facilitate integration across experiments

We therefore examined the antibody-derived tags (ADT) for 173 cell surface proteins, measured simultaneously in each experiment (Supplementary Figure 1C). For example, in the H3K4me1 dataset, ADT expression patterns were fully concordant with cell types determined from chromatin-based clustering (i.e. uniform expression of CD3 in T cells, CD14 in monocytes, and CD19 on B cells; Supplementary Figure 1D). However, the protein modality revealed additional substructure, including mutual exclusivity of CD4 and CD8 in T cells, a division of monocytes into classical CD14+ and nonclassical CD16+ subgroups, immunoglobulin heterogeneity correlating with B cell maturation, and canonical marker expression denoting Mucosal Associated Invariant T (MAIT) (Figure 1F, Supplementary Figure 1D). We utilized our recently introduced Weighted Nearest Neighbor (WNN) analysis^30^ (Supplementary Methods) to simultaneously cluster scCUT&Tag-pro based on a weighted combination of modalities (Figure 1F). For each experiment, we found that the protein modality substantially enhanced our ability to separate distinct cell types, echoing our previous findings in CITE-seq and ASAP-seq analysis^26,28^ (Supplementary Figure 2A).

We compared the results of our single-cell analysis with FACS-sorted H3K4me1 ChIP-seq profiles from ENCODE^35^ (Supplementary Methods). We observed that pseudobulk profiles generated from our WNN-derived scCUT&Tag-pro clusters clearly recapitulated the cell type-specific ENCODE profiles (Figure 1G). Moreover, downsampling our single-cell clusters to as low as 300 cells/profile still retained substantial cell type-specificity in the ensuing pseudobulk tracks (Figure 1G). Taken together, these analyses demonstrate that scCUT&Tag-pro datasets are comparable in sensitivity to existing technologies, exhibit low background, contain sufficient information to identify granular cell types and states, and can be used to generate pseudobulk profiles that reflect high-quality bulk data when a sufficient number of cells are included.

As an alternative to fully unsupervised analysis, we found that single-cell protein measurements also facilitated high-resolution supervised annotation using reference datasets. For example, we recently introduced a carefully annotated reference dataset of 58 major and minor cell types and states in the circulating human immune system, using a CITE-seq dataset of 161,764 cells and 224 surface proteins^30^. As our protein panel in this study largely overlaps with the reference dataset, we ‘mapped’ the cells from each scCUT&Tag-pro experiment onto our reference dataset (Supplementary Methods), and repeated the procedure for our published ASAP-seq dataset^28^ as well. Performing query-to-reference mapping using shared protein markers enables joint visualization of all datasets (Figure 2A and 2B), and also transfers a consistent set of annotations to each cell at multiple levels of resolution. This enabled us to partition each dataset into broad (level 1), granular (level 2), and fine-grained (level 3) classifications (Supplementary Figure 2B). We also utilized this approach to infer a unified pseudotime trajectory (Supplementary Methods) that modeled CD8 T cell transitions from naive to effector states and confirmed the accuracy and resolution of our trajectory by comparing the developmental dynamics of key markers across experiments (Figure 2C, Supplementary Figure 2C).

**Figure 2:**
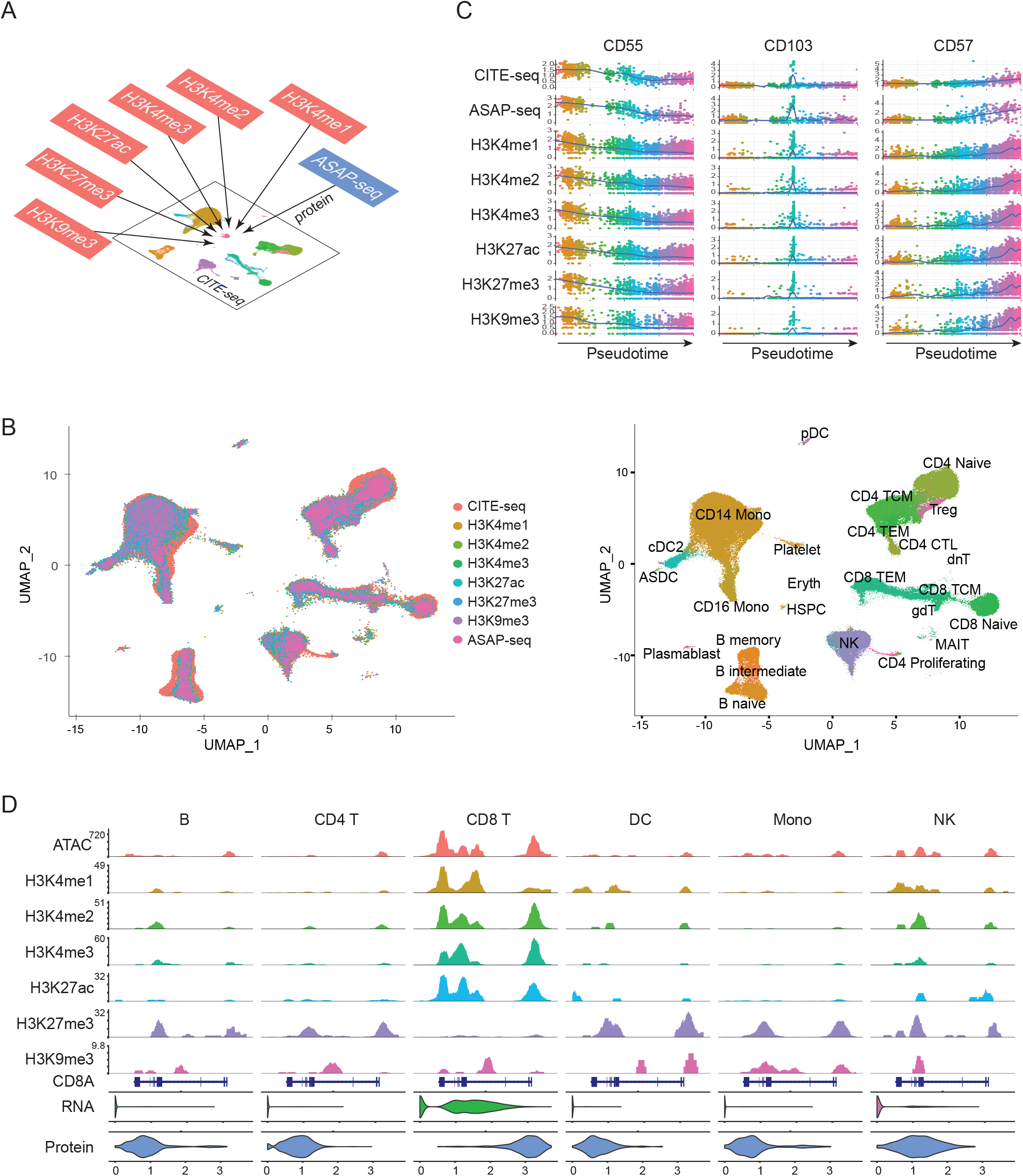
Protein measurements facilitate integrated analysis across modalities. **(A)** Schematic workflow for integrated analysis. Datasets produced by scCUT&Tag-pro, ASAP-seq, and CITE-seq are integrated together on the basis of a shared panel of cell surface protein measurements. **(B)** Left: UMAP visualization of 230,597 total cells projected onto the reference dataset from (Hao et al, Cell 2021). Right: In addition to a harmonized visualization, cells from all experiments are annotated with a unified set of labels. **(C)** We learned a unified pseudotime trajectory based on all experiments, representing the CD8 T cell transition from naive to effector states. We observe identical molecular dynamics for naive (CD55), memory (CD103), and effector (CD57) markers across all experiments, demonstrating that integrative analysis accurately identifies cells in matched biological states across experiments. **(D)** Visualization of nine molecular modalities at the CD8A locus in B cell (B), CD4 T cell (CD4 T), CD8 T cell (CD8 T), dendritic cell (DC), monocyte (Mono) and natural killer cell (NK) groups.

Our reference-mapping workflow enables us to explore the relationships between nine different modalities by harmonizing them into a common space and providing a consistent set of annotations (Figure 2B). The modalities span the central dogma and range from measurements of chromatin accessibility, protein-DNA interactions (six targets), gene expression, and protein abundances. While the individual modalities are collected in separate experiments, each originates from a multimodal technology (ASAP-seq, scCUT&Tag-pro, CITE-seq), that is paired with a large and comprehensive panel of surface protein measurements to facilitate integration. For example, in Figure 2D we visualize data from each modality collected at the CD8A locus. As expected, we find robust gene and protein expression (as measured by CITE-seq), enriched chromatin accessibility (as measured by ASAP-seq), and the presence of activating histone modifications (as measured by scCUT&Tag-pro) in CD8 T cells. In myeloid cell types, the locus is characterized by inaccessible chromatin and the presence of repressive histone modifications, which results in undetectable expression of CD8 RNA and protein.

### scChromHMM characterizes chromatin state at single-cell resolution

We next aimed to leverage our dataset to explore heterogeneity in chromatin state within the human immune system. We first considered an analysis strategy exclusively leveraging tools that have been previously developed for bulk chromatin analysis and provide 25 pseudobulk CUT&Tag tracks (based on level-2 reference derived annotations) as input. For example, ChromHMM^12,17^ trains a hidden markov model (HMM) on the concatenated histone modification profiles for all cell types, and applies the Baum-Welch training algorithm^36^ to identify a set of possible hidden ‘chromatin states’, each of which corresponds to a coherent combination of histone marks. ChromHMM then applies the forward-backward algorithm^36^ individually on each pseudobulk track, in order to calculate the posterior probability that each 200bp genomic window in each cell type is assigned to a particular chromatin state.

After running the Baum-Welch step (Supplementary Methods), ChromHMM obtained 12 states (Figure 3A), which could be broadly grouped into active promoter (enriched for H3K4me3/H3K4me2/H3K27ac), active enhancer (enriched for H3K4me1/H3K27ac), repressed (enriched for H3K27me3), and heterochromatic (enriched for H3K9me3) states. We also observed more subtle heterogeneity in chromatin state within these broad categories, likely corresponding to strong/weak states as has been previously described^17^. These results suggest that the Baum-Welch step of ChromHMM can be successfully run on pseudobulk tracks generated from scCUT&Tag-pro, returning a functionally interpretable set of chromatin states.

**Figure 3:**
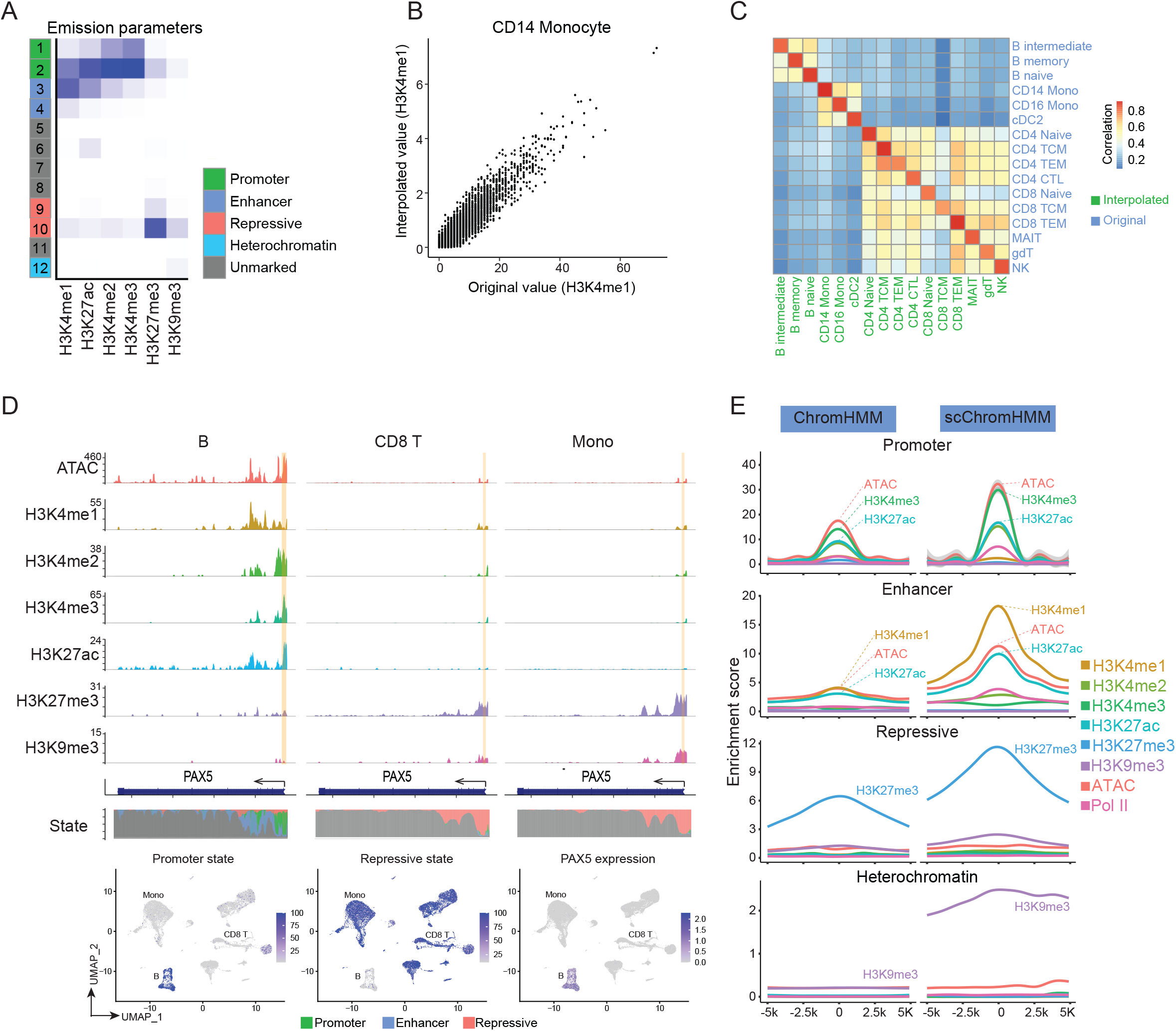
scChromHMM annotates chromatin states at single-cell resolution. **(A)** Chromatin states returned by ChromHMM, which was run on 25 pseudobulk tracks for six histone marks. States can be broadly grouped into five categories. **(B)** Correlation comparing cell type-specific pseudobulk profiles of H3K4me1 in CD14 monocytes, generated from the original experiment, or from the interpolated values. Each point corresponds to a 200bp genomic window (Supplementary Methods) **(C)** For each cell type, the interpolated and original profiles are highly correlated and clustered together. **(D)** scChromHMM outputs at the PAX5 locus. (Top) Pseudobulk profiles for six chromatin marks in three cell types. Yellow bar highlights a 200bp genomic window near the TSS. (Bottom) scChromHMM posterior probabilities representing the annotation for the highlighted window in each cell. The region is uniformly annotated with a promoter state in B cells where PAX5 is transcriptionally active, and as a repressive state in other cell types. **(E)** Metaplots exhibiting the enrichment of chromatin accessibility and histone modifications at functional regions identified by chromHMM (left) and scChromHMM (right) in CD14 monocytes.

However, when attempting to associate these states with individual genomic elements in each cell type, we identified a limitation associated with the application of bulk methods to single-cell datasets. In particular, we found that for this step, the results of ChromHMM were highly dependent on the level of granularity used when generating pseudobulk profiles. For example, when we considered genomic windows assigned to enhancer states in CD8+ T cells (level 1), 16.4% were not annotated as enhancers in any of the more granular T cell subsets when we repeated the analysis using a higher-resolution subset of cell labels (level 2). Especially when analyzing developmental tissues, or systems characterized by continuous sources of heterogeneity, it may be detrimental to condition all downstream analyses on a fixed set of discrete cell type labels^37^.

We therefore devised an alternative approach, scChromHMM, where chromatin states are assigned at single-cell resolution. In order to do so, we first generate single-cell profiles with simultaneous measurements of six histone marks. While it is not currently feasible to experimentally generate these data, we leveraged our previously described anchoring workflow^38^ as a computational alternative. We recently demonstrated the use of this workflow to ‘transfer’ modalities across experiments^38^, for example, using a CITE-seq reference to accurately and robustly predict cell surface protein levels in an scRNA-seq dataset of human bone marrow. We apply a similar procedure in this study (Supplementary Methods), in order to interpolate 20,000 single-cell profiles, each of which consists of quantitative and genome-wide profiles for all six histone modifications and chromatin accessibility, alongside RNA and protein measurements.

We carefully assessed the accuracy of our interpolated profiles by comparing them to the original measurements that were obtained in separate experiments. For example, we computed the pseudobulk profile for H3K4me1 measurements based on our interpolated profiles, or the original measured values, and observed high quantitative concordance (Figure 3B; R=0.95). For each cell type, pseudobulk profiles of interpolated measurements clustered specifically with the pseudobulk profiles from the original measurements (Figure 3C, additional histone marks shown in Supplementary Figure 3A), demonstrating that our procedure retains cell type-specific variation. Finally, we repeated the interpolation procedure to generate an independent set of 20,000 profiles and observed high reproducibility (R=0.98) between runs (Supplementary Figure 3B). While our procedure cannot capture stochastic fluctuations, it does represent a form of data ‘denoising’ by averaging histone modification signal across cells in similar biological states, and therefore alleviates the sparsity limitations associated with scCUT&Tag measurements. We emphasize that the acquisition of protein measurements is essential for this strategy, as the protein modality enables the accurate identification of anchors across diverse datasets.

Having acquired interpolated chromatin profiles for six histone modifications at single-cell resolution, we next aimed to annotate the chromatin state of each 200bp genomic window, in each cell. To achieve this, we ran the forward-backward algorithm^12,36^ individually on each cell (Supplementary Methods), using our interpolated profiles for six histone modifications and our previously calculated set of emission and transition probabilities as input. We note that data binarization is a requirement for this procedure and found that the frequency of our interpolated profiles across genomic windows exhibited clear evidence of bimodality, facilitating this process (Supplementary figure 4A). The output of scChromHMM represents, for each genomic window in each cell, the posterior probability distribution across 12 chromatin states. For example, in Figure 3D, we visualize the posterior probabilities (promoter) of a genomic window located at the PAX5 transcriptional start site (TSS), at single-cell resolution.

Regions classified by scChromHMM as active promoters were enriched near the TSS of actively transcribed genes, active enhancer regions were distal to the TSS but enriched for accessible chromatin, and heterochromatic regions largely overlapped (76.3%) with a set of repetitive elements annotated by RepeatMasker^39^ (Figure 3E, Supplementary Methods). We also found that putatively functional regions identified by scChromHMM exhibited increased accuracy and robustness compared to regions identified by ChomHMM run on pseudobulk profiles. Regions classified by scChromHMM consistently exhibited stronger enrichment for key histone modifications (Figure 3E). Additionally, we performed a scCUT&Tag-pro experiment to measure binding for RNA Polymerase II (Ser2/Ser5 phosphorylated), which was withheld from chromatin state prediction. Regions annotated as active promoters by scChromHMM exhibited greater enrichment for RNA PolII compared to active promoters identified by ChromHMM (Supplementary Figure 4B). We conclude that scChromHMM enables chromatin state predictions at single-cell resolution, and also improves the accuracy of state prediction.

### Heterogeneity in repressive chromatin state suggests regulatory priming

We utilized the output of scChromHMM to explore the dynamics of chromatin state changes across a trajectory of CD8 T cell maturation from naive to effector states. For example, we identified 14,585 genomic windows that acquired or lost a repressive chromatin state (Supplementary Methods), indicating that extensive remodeling of repressive chromatin is associated with this cellular process, particularly at the sharp transition from naive to memory cells (Figure 4A). In order to explore motifs that were associated with these transitions, we modified the chromVAR algorithm^40^. While chromVAR traditionally identifies DNA motifs whose presence is associated with increased genome-wide chromatin accessibility, we instead searched for motifs whose presence was correlated with an increased posterior probability across the learned chromatin states (Supplementary Methods).

**Figure 4:**
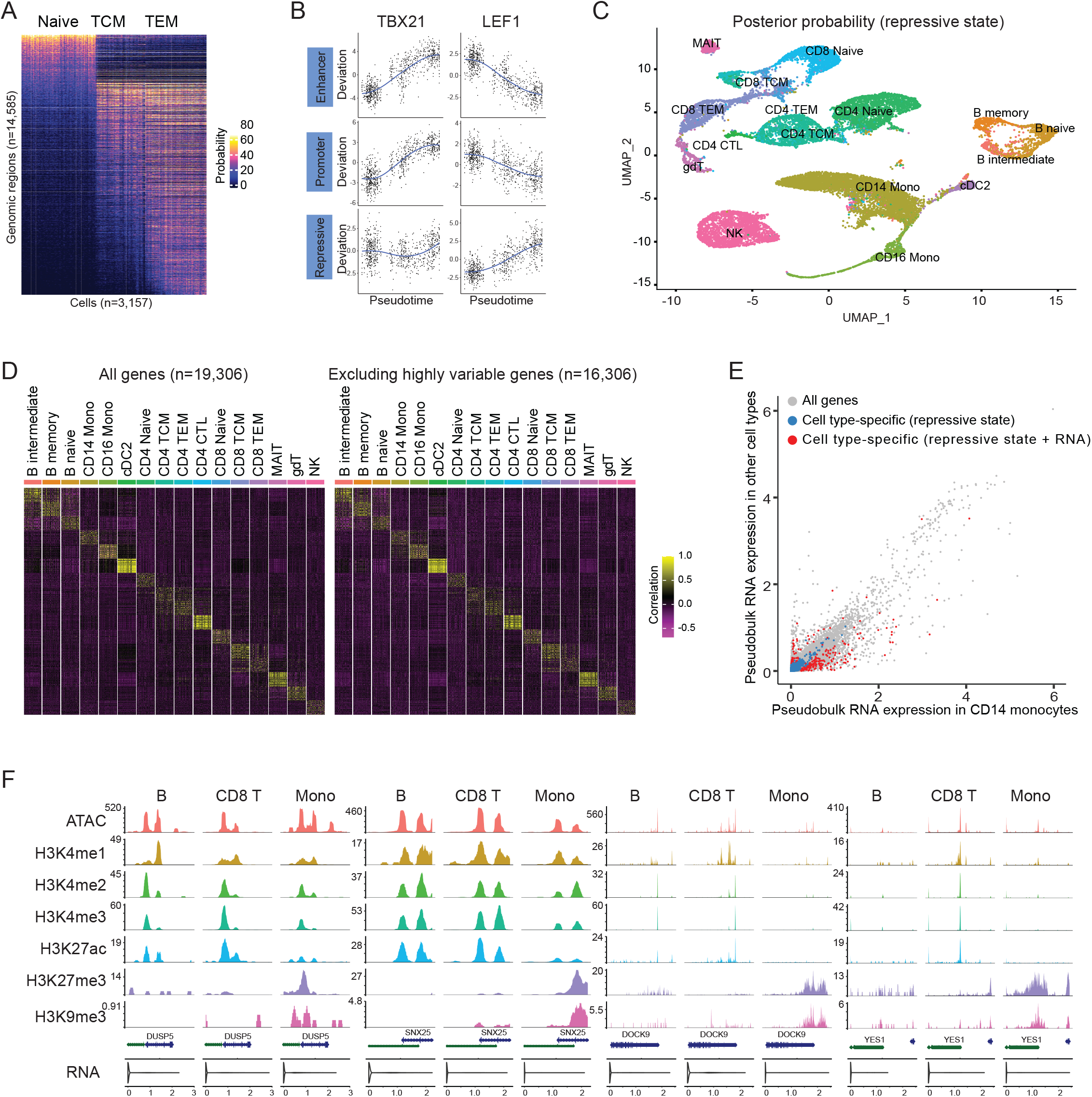
Extensive heterogeneity in repressive chromatin encodes cellular identity. **(A)** Remodeling of repressive chromatin during CD8 T cell maturation. Heatmap shows the posterior probabilities (repressive state) in single cells for 14,585 genomic loci, as returned by scChromHMM. Cells are ordered by their progression along pseudotime (Figure 2C). **(B)** ChromVar deviation scores for the TBX21 and LEF1 motifs in single cells, ordered by their progression along pseudotime. We used the scChromHMM-derived posterior probabilities as input to ChromVar, instead of chromatin accessibility levels. **(C)** Unsupervised analysis of scChromHMM-derived probabilities (repressive state) separates granular cell types. **(D)** Single-cell correlation matrix based on repressive chromatin at TSS (Supplementary Methods) when using all TSS (left heatmap), or after excluding the top 3,000 transcriptionally variable genes (right heatmap). In each case, the observed correlation structure is fully consistent with cell type labels, suggesting that there is extensive heterogeneity in repressive chromatin even for genes that do not vary transcriptionally.(**E)** Scatter plot showing average gene expression levels for all genes in CD14 monocytes (x-axis) and other cell types (y-axis). Colored points represent 1,597 loci where we detect changes in repressive chromatin for monocytes (Supplementary Methods). Blue points represent 1,340 loci where we do not detect an accompanying transcriptional change. Red points represent 257 genes where we detect a transcriptional shift. **(F)** Four representative examples of individual genes shown as blue points in (E).

Our analyses revealed that genomic windows containing the motif bound by the transcription factor TBX21 (T-bet), which is a crucial regulator of T cell responses^41,42^, exhibit highly dynamic chromatin states during T cell maturation (Figure 4B). In particular, as cells transition from naive to memory and effector states, genomic windows containing TBX21 motifs acquire an increased probability of entering an active promoter or enhancer state. Conversely, genomic windows containing motifs bound by the critical T cell regulator LEF1^43^ tend to exist in active chromatin states at the onset of the trajectory, but the same windows transition to acquire a repressive state as cells progress through the trajectory (Figure 4B). More generally, when aggregating cells into granular subgroups, we observed substantial cellular heterogeneity and cell type-specificity for motif activity scores computed based on scChromHMM-determined probabilities (Supplementary Figure 4C). This analysis nominated a suite of regulators for each cell type that were also differentially expressed at the transcriptional level (Supplementary Figure 4C). Taken together, our results demonstrate the potential for integrated analyses of scCUT&Tag data to identify DNA sequence motifs that correlate with diverse chromatin states.

We next asked if cellular heterogeneity in repressive chromatin state was fully reflected in the cellular transcriptome. We first considered the scChromHMM derived probabilities for the repressive state at genomic windows overlapping TSS. We found that granular cellular states could be separated based on unsupervised clustering of these measurements (Supplementary Methods), indicating that cellular identity is encoded not only in transcriptional output or immunophenotypes, but also in genome-wide profiles of repressive chromatin state (Figure 4C). To our surprise, we found that granular cellular identity remained encoded in repressive chromatin probabilities at TSS, even after excluding the 3,000 most variable genes in the transcriptome (Figure 4D). This suggested that even for genes whose expression did not vary across cell types, there may be substantial heterogeneity in chromatin state.

We therefore considered the set of loci where we observed cell type-specific changes in repressive chromatin state (Supplementary Methods) and calculated their overlap with transcriptional changes as well. For example, we observed 1,597 TSS loci whose posterior probability (repressive chromatin state) and H3K27me3 signal was enriched or depleted in CD14 monocytes. Within these associated genes for these loci, we detected monocyte-specific transcriptional shifts in only 257 (16.1%) (Figure 4E and 4F). When we did observe both transcriptional and chromatin-based shifts, they were highly concordant (i.e. the acquisition of a repressive state was associated with a decrease in gene expression). However, when we identified heterogeneity in repressive chromatin state in the absence of transcriptional variation, gene expression was either absent or at very low levels in all cell types (Figure 4E). When considering their pseudobulk expression, the median expression of the set of 1,340 genes in PBMC was below 0.5 transcripts per million. These genes are not informative when determining cellular state from scRNA-seq data, yet their repressive chromatin landscape exhibits clear patterns of cell type-specificity (Figure 4F).

Gene Ontology ontology analysis^44^ of these gene sets was enriched for DNA-binding transcription factors, and transmembrane and channel proteins whose activity may regulate the activation of downstream signaling pathways and immune responses (Supplementary Figure 4D). We hypothesize that these chromatin shifts may represent a form of regulatory ‘poising’, that is not yet reflected in the current transcriptional output of each cell but may be relevant to its future behavior or potential. Consistent with this hypothesis, a recent study of H3K27me3 heterogeneity profiled in bulk samples indicated that variability in repressive chromatin represented a signature for genes involved in establishing and regulating cell state^45^. We note that some genes exhibiting poised behavior may in fact be differentially expressed across cell types but cannot be accurately quantified by scRNA-seq.

### Projecting additional scCUT&Tag datasets onto a multimodal atlas

Our analyses demonstrate the utility of projecting data from diverse molecular modalities into a harmonized manifold, creating a multimodal atlas that spans CITE-seq, ASAP-seq, and scCUT&Tag-pro experiments. While the construction of this multimodal reference required the presence of protein levels in each experiment, it is not always possible to measure cell surface protein levels when performing scCUT&Tag, particularly when performing analysis of single nuclei. We therefore considered how to project unimodal scCUT&Tag profiles into our reference dataset, even in the absence of accompanying protein measurements.

We have recently released a framework, Azimuth^30^, which addresses a similar problem of projecting scRNA-seq datasets onto a CITE-seq defined human PBMC reference. In Azimuth, we leverage a semi-supervised dimensionality reduction approach, supervised PCA^46^, which identifies the best transcriptomically-defined vectors that separate CITE-seq defined cell types. In our previous work we have shown that supervised PCA outperforms unsupervised dimensional reduction approaches when projecting new scRNA-seq datasets onto a multimodal reference^30^. We pursued a similar strategy to map a recently published H3K27me3 scCUT&Tag dataset of human PBMC nuclei^19^ onto our multimodal reference atlas. Using our scCUT&Tag-pro dataset as a reference, we calculated a modified LSI (Supplementary Methods) which transforms the histone modification profiles into a low-dimensional space that retains separation of our multimodally defined cell types. We utilize this transformation to identify anchors and ‘map’ the query dataset into our reference dataset (Supplementary Methods).

We observe that by performing reference-mapping we could substantially improve the interpretation of the query dataset, and in particular, the ability to resolve granular cell types. For example, unsupervised clustering of the query dataset^19^ was consistent with the previously published analysis and revealed broad clusters of immune subsets, including Monocytes, B, and T/NK subroups (Figure 5A). After reference mapping, monocytes could be segregated into CD14 and CD16+ subsets, T/NK subclusters could be further resolved into CD4 T, CD8 T, and NK groups with additional heterogeneity for naive and effector states, and B cells separated into different developmental stages (Figure 5B). We found that we could classify (prediction score >0.5) 88% of cells at level-1 resolution, and 73% of cells at level-2 resolution. We find high concordance when comparing patterns of cell type-specific H3K27me3 binding compared in both the reference and query datasets (Figure 5C-E), verifying the accuracy of our predictions. Therefore, we propose that our multimodal atlas represents a broad scaffold for the circulating human immune system that can be used to map datasets from a wide variety of cutting-edge single-cell technologies.

**Figure 5:**
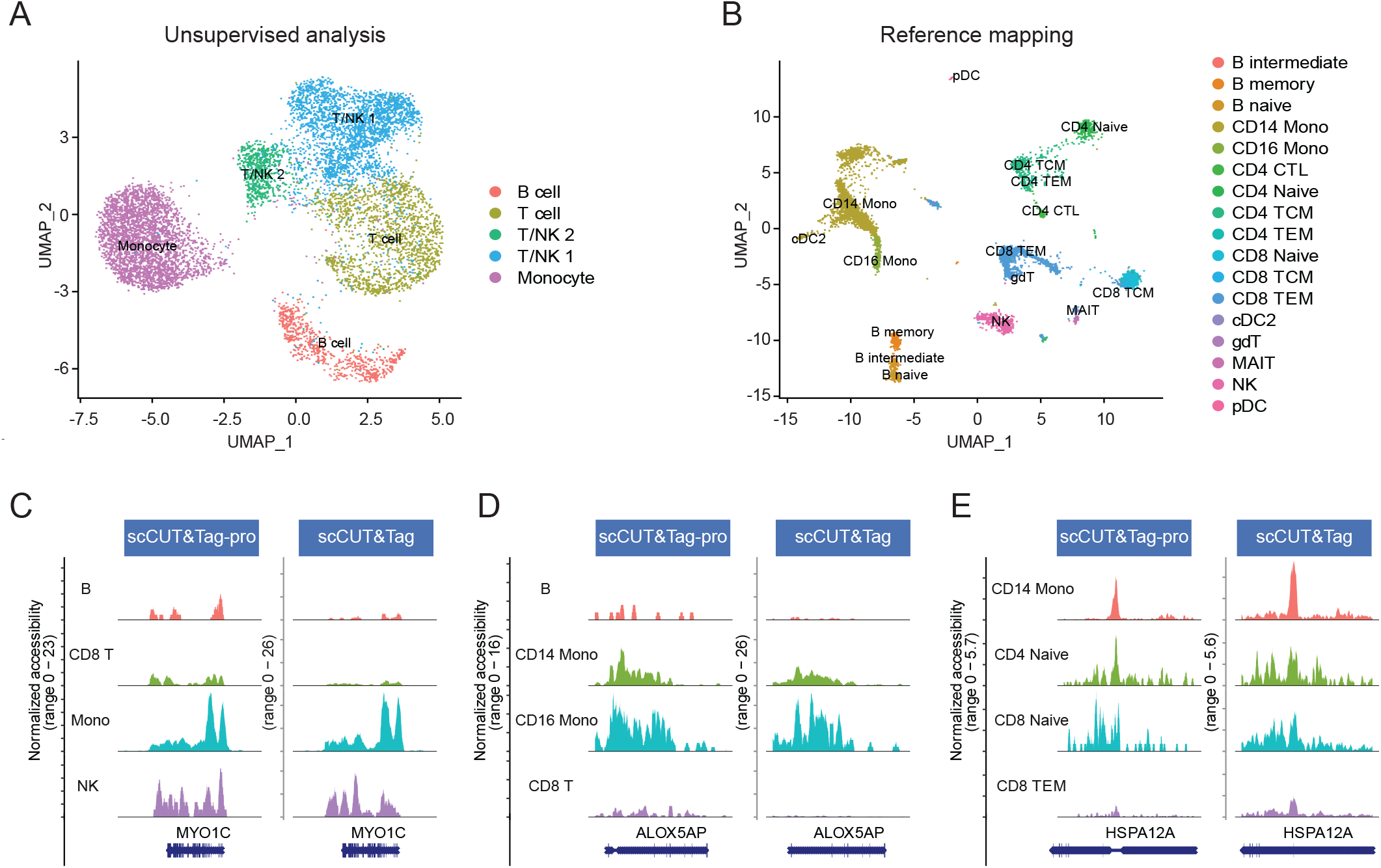
Supervised mapping of scCUT&Tag datasets. **(A)**UMAP visualization of 8,536 H3K27me3 scCUT&Tag profiles of human PBMC, from (Wu et al, 2021), based on an unsupervised analysis and clustering. **(B)** Same cells as in (A), but after mapping to the multimodal reference defined in this paper. Cells are colored by their reference-derived level 2 annotations. **(C)-(E)** Coverage plots showing the cell type-specific binding patterns of H3K27me3 at three loci. Plots are shown for our dataset (reference), as well as the scCUT&Tag profiles from the query dataset (query, Wu et al, 2021). Cells in the query dataset are grouped by their predicted labels. We observe highly concordant patterns across datasets for all loci, supporting the accuracy of our predictions. Four representative cell types are shown at each locus. TEM: T effector memory.

## Discussion

One of the key goals of multi-omic single-cell studies is to explore the relationships across different molecular modalities, and to explore how changes in one modality affect variation in another. Here, we pursue an integrative analysis where nine different molecular modalities are each measured in separate experiments. We subsequently integrate these data and derive what we colloquially refer to as single-cell ‘mega-omic’ profiles, each representing an individual cell containing measured or interpolated measurements for many modalities. In addition, we introduce scChromHMM, which enables the exploration of heterogeneity in chromatin state across both discrete cell types and continuous trajectories.

Our integrated analysis strategy represents an analytical alternative to technological solutions that aim to simultaneously profile multiple modalities in single cells. Despite substantial recent progress^22,23^, it is not currently feasible to simultaneously profile the genome-wide binding patterns of six different histone modifications in the same cell, particularly when there is a partial overlap in their localization. Similarly, future extensions of CUT&Tag enabling simultaneous measurements of RNA and protein levels are likely to result in decreased per-modality data quality, as has been previously described^28,29^. Our approach therefore represents a feasible and broadly applicable alternative, and we demonstrate that our ‘mega-omic’ profiles are highly quantitative, sensitive, and robust. However, interpolated modality predictions cannot capture stochastic technical or biological variation or be used to detect associations between multiple histone modifications within the same cell.

The successful integration of CITE-seq, ASAP-seq, and scCUT&Tag-pro datasets originates from a shared panel of cell surface protein levels that are collected in each experiment. We emphasize that cell surface protein levels represent one of multiple potential modalities that can be utilized for ‘mega-omics’ integration. For example, two recent pioneering approaches (Paired-Tag^22^, and CoTECH^23^) introduced combinatorial indexing-based approaches for simultaneous scCUT&Tag and scRNA-seq in single cells, enabling the integration of additional multi-omic technologies (i.e. Paired-Seq) via transcriptomic measurements. Our scCUT&Tag-pro technology represents a complementary solution that is compatible with the 10x Genomics Chromium system, and will be particularly valuable for profiling immune cell types that are well-defined by their surface protein landscape, yet remain challenging to profile via combinatorial-indexing.

To our surprise, we found substantial heterogeneity in chromatin state even at genes that did not exhibit transcriptional variation. In particular, we observed cell type-specific repression, as demarcated by the presence of H3K27me3, even for genes that were expressed at similarly low levels across all cell types. While it is possible that this heterogeneity in repressive chromatin state has minimal functional consequence, cellular variation in chromatin state may also underlie future cell type-specific transcriptional responses. For example, Mellis et al. recently found that cell type specificity was encoded not only in steady-state transcriptional output, but also in future responses to environmental perturbations^45^. Similarly, Shim et al. identified that genes with heterogeneous H3K27me3 landscapes across distinct organ systems were strikingly enriched for key developmental, regulatory, and morphological functions^47^. We propose that scCUT&Tag-pro represents a sensitive and powerful technique for identifying loci exhibiting heterogeneous repressive chromatin within an organ system as well. Future experiments profiling chromatin state in the bone marrow will help to elucidate the developmental dynamics that establish this heterogeneity.

Moving forward, we expect that integrated analyses across multiple modalities will not only help to understand the molecular state of single cells, but also will reveal new insights on fundamental questions in gene regulation. For example, the activation of gene expression is correlated with changes in chromatin accessibility, the acquisition and loss of histone modifications, the binding of transcription factors and RNA polymerase, and the formation of new DNA contacts. However, the regulatory relationships and temporal ordering between these steps remains poorly understood. Motivated by the ability of RNA velocity analysis to leverage known temporal relationships between unspliced and spliced reads in order to infer the direction of developmental trajectories^48^, we envision that single-cell multi-omic analysis will help to order the coordinated molecular events that collectively establish cellular heterogeneity.

## Data availability

Seurat, Signac, and scChromHMM are freely available as open-source software packages:

https://github.com/satijalab/seurat

https://github.com/timoast/signac

https://github.com/satijalab/scchromhmm

The scCUT&Tag-pro datasets are available as open-access downloads at: https://zenodo.org/record/5504061

## Acknowledgements

The authors would like to thank all the members of the Satija Lab for thoughtful discussions related to this work. B.Z. is a postdoctoral fellow of the Jane Coffin Childs Memorial Fund for Medical Research. This investigation has been aided by a grant from the Jane Coffin Childs Memorial Fund for Medical Research. This work was supported by the Chan Zuckerberg Initiative (EOSS-0000000082, HCA-A-1704-01895 to R.S.), and the NIH (K99HG011489-01 to T.S; RM1HG011014-02, 1OT2OD026673-01, DP2HG009623-01 to R.S).

## Supplementary Methods

### PBMC acquisition and processing

Cryopreserved healthy donor PBMCs were purchased from AllCells. After thawing into DMEM with 10% FBS, the cells were spun down at 4°C for 5 min at 400 g and washed twice with PBS with 2% BSA. After centrifugation, the cell pellet is resuspended in staining buffer (2% BSA and 0.01% Tween in PBS).

### scCUT&Tag antibodies

The antibodies used were H3K4me1 (1:100, Abcam, ab8895), H3K4me2 (1:100, Abcam, ab32356), H3K4me3 (1:100, Abcam, ab213224), H3K27ac (1:100, Abcam, ab177178), H3K27me3 (1:100, Cell Signaling Technology, 9733), H3K9me3 (1:100, Abcam, ab8898), Phospho-Rpb1 CTD (Ser2/Ser5) (1:50, Cell Signaling, 13546) and guinea pig anti-rabbit (1:100, Novus Biologicals, NBP1-72763). TotalSeq-A conjugated antibodies and panels were obtained from BioLegend (399907, see Supplementary Table 2 for a list of antibodies, clones and barcodes).

### scCUT&Tag-pro experimental workflow

#### Surface Protein Antibody staining

5 million thawed PBMC were resuspended in 200 *µ*L staining buffer (2% BSA and 0.01% Tween in PBS) and incubated for 15 min with 10 *µ*L Fc receptor block (TruStain FcX, BioLegend) on ice. The samples were evenly distributed into 5 tubes and 2 *µ*L hashing antibody was added separately. Cells were then washed three times with 1 mL staining buffer and pooled together. 2 million PBMC were used for each histone modification. After cell hashing, the panel of oligo-conjguated antibodies (one test is sufficient for 2 million cells) was added to the cells to incubate for 30 min on ice. After staining, cells were washed three times with 1 mL staining buffer and resuspended in 100 *µ*L staining buffer. Subsequently, 4 *µ*L Fab Fragment Goat Anti-Mouse IgG (Jackson ImmunoResearch, 115-007-003) was added for incubation on ice for 15 min. The cells were then washed three times with 1 mL staining buffer. After the final wash, cells were resuspended 200 μl PBS ready for fixation.

#### Fixation and permeabilization

1.25 *µ*L of 16% Methanol-free formaldehyde (Thermo Fisher Scientific, PI28906) was added for fixation (final concentration: 0.1%) at room temperature for 5 min. The cross-linking reaction was stopped by addition of 12 *µ*L 1.25M glycine solution. Subsequently, cells were washed twice with PBS. The permeabilization was performed by adding isotonic lysis buffer (20 mM Tris-HCl pH 7.4, 150 mM NaCl, 3 mM MgCl2, 0.1% NP-40, 0.1% Tween-20, 1% BSA, 1× protease inhibitors) on ice for 7 min. Subsequently, 1 mL of cold wash buffer (20 mM HEPES pH 7.6, 150 mM NaCl, 0.5 mM spermidine, 1× protease inhibitors) was added, and cells were centrifuged at 800g for 5 min at 4°C.

#### Tagmentation

Permeabilized cells were directly resuspended with 150 *µ*L antibody buffer (20 mM HEPES pH 7.6, 150 mM NaCl, 2 mM EDTA, 0.5 mM spermidine, 1% BSA, 1× protease inhibitors) with primary antibody (for example, anti-H3K4me1) and incubated overnight on a rotator at 4°C. The next day, cells were washed once with 150 μl wash buffer and centrifuged for 5 min at 800g. After removing the supernatant, the cells were resuspended in 150 μL of wash buffer with secondary antibody and incubated for 1h at room temperature on rotator. The cells were washed three times with 150 μL wash buffer to remove excess remaining antibodies. The cells were then resuspended in 150 μL high salt wash buffer (20 mM HEPES pH 7.6, 300 mM NaCl, 0.5 mM spermidine, 1× protease inhibitors) with 7.5 *µ*L pAG-Tn5 (EpiCyper, 15-1017) and incubated for 1 h on rotator at room temperature. The cells were then washed twice with high salt wash buffer and resuspended in 100 μL tagmentation buffer (20 mM HEPES pH 7.6, 300 mM NaCl, 0.5 mM spermidine, 10 mM MgCl2, 1× protease inhibitors). The samples were placed on PCR machine to incubate for 1h at 37 °C. When tagmentation was done, the reaction was stopped by adding 4 μL of 0.5 M EDTA. Tagmentation steps were performed in 0.2 mL tubes in order to minimize cell loss.

#### Single-cell encapsulation, PCR, and library construction

After tagmentation, cells were centrifuged for 5 min at 1000g and the supernatant was discarded. Cells were resuspended with 30 *µ*L 1× Diluted Nuclei Buffer (10x Genomics), counted, and diluted to a concentration based on the targeted cell number. The transposed cell mix was prepared as following: 7 *µ*L of ATAC buffer and 8 *µ*L cells in 1× Diluted Nuclei Buffer. All remaining steps were performed according to the 10x Chromium single-cell ATAC protocol and the library construction method was adapted from ASAP-seq^28^. Briefly, 0.5 μL of 1 μM bridge oligo A (TCGTCGGCAGCGTCAGATGTGTATAAGAGACAGNNNNNNNNNVTTTTTTTTTTTTTTTTTTTT TTTTTTTTTT/3InvdT/) was added to the barcoding mix. Linear amplification was performing using the following PCR program: (40 °C for 5 min, 72 °C for 5 min, 98 °C for 30 s; 12 cycles of 98 °C for 10 s, 59 °C for 30 s and 72 °C for 1 min; ending with hold at 15 °C). The remaining steps were performed according to the 10x scATAC-seq protocol (v1.1), with the following additional modifications:

Antibody-derived tags: During silane bead elution (Step 3.1s), beads were eluted in 43.5 μL of elution solution I. The extra 3 μL was used for the surface protein tags library. During SPRI cleanup (Step 3.2d), the supernatant was saved and the short DNA derived from antibody oligos was purified with 2.0× SPRI beads. The eluted DNA was combined with the 3 *µ*L left aside after the silane purification to be used as input for protein tag amplification. PCR reactions were set up to generate the protein tag library with KAPA Hifi Master Mix (P5 and RPI-x primers): 95 °C for 3 min; 14–16 cycles of 95 °C for 20 s, 60 °C for 30 s and 72 °C for 20 s; followed by 72 °C for 5 min and ending with hold at 4 °C.

P5 primer: AATGATACGGCGACCACCGAGATCTACAC RPI-x primer: CAAGCAGAAGACGGCATACGAGATxxxxxxxxGTGACTGGAGTTCCTTGGCACCCGAGAATTC CA.

#### Sequencing

The final libraries were sequenced on NextSeq 550 with the following recipe: i5:16bp, i7: 8bp, read1: 34bp, read2: 34bp.

### scCUT&Tag-pro data preprocessing

#### Antibody-derived tags and Cell Hashing libraries

We use salmon alevin 1.4.0^49^ for the alignment and quantification of single-cell antibody-derived tags (ADT) and hashtag oligos (HTO), using a custom index that is constructed only from known ADT and HTO barcode sequences. We apply a threshold of 10 protein and 10 hashing antibody counts. Cells that pass these thresholds are used as input to the HTODemux() function in Seurat^30^ for demultiplexing and doublet removal. Antibody data was normalized using the centered log ratio transformation. In order to visualize cells on the basis of their ADT profiles, we performed PCA using all features as input, and retained the first 25 principal components to construct a nearest-neighbor graph and perform UMAP visualization.

#### pAG-Tn5 tagmented libraries (i.e. scCUT&Tag)

We use cellranger-atac v1.2 pipeline with default settings to align reads to the hg38 human reference annotation and generate fragment files. We filter the fragment file to keep the cellular barcodes (CBs) that were present in the filtered ADT and HTO libraries, and also contained at least 100 fragments. We use the AggregateTiles function from the Signac package^50^ to quantify the fragment coverage based on 5,000bp genomic windows, remove windows with less than 5 counts, and then merge adjacent windows. We next perform modified TD-IDF normalization^50^, which corrects for differences in cellular sequencing depth and weights each window based on its average fragment abundance across all cells. We next run singular value decomposition (SVD) on the TF-IDF matrix to generate the low dimensional representation of the data. We use dimensions 2:30 for UMAP visualization, as the first dimensional is highly correlated with sequencing depth.

We utilize the Seurat v4 Weighted Nearest Neighbor (WNN) framework^30^ to generate a multimodal representation of the scCUT&Tag-pro datasets. We use the FindMultiModalNeighbors function to generate a WNN graph, using dimensions 1:25 (ADT modality) and dimensions 2:30 (scCUT&Tag modality).

#### Reference mapping

We integrate each of our scCUT&Tag-pro datasets with a previously published reference CITE-seq dataset^30^. In previous work, we apply ‘supervised PCA’ (sPCA) to identify a linear projection of the scRNA-seq measurements in the CITE-seq dataset that maximally retains the information captured in both modalities. This transformation can be applied to new scRNA-seq query datasets in order to identify anchors with the CITE-seq reference. These anchors are used for annotation and visualization of the query cells. Our framework for query-to-reference mapping is implemented as part of our Azimuth tool, accessible at azimuth.hubmapconsortium.org and fully described in our previous paper^30^.

Here, we leverage a similar strategy, but perform sPCA on the protein measurements from the CITE-seq dataset. As input to sPCA, we use all 128 protein features that were measured both in this study as well as in the reference dataset. This procedure identifies a linear transformation of cell surface protein measurements that can be applied to scCUT&Tag-pro measurements. We use the FindTransferAnchors anchors (reduction = ‘spca’, dims = 1:30) in Seurat v4 to identify anchors between reference and query datasets, and the MapQuery function (default parameters) in order to annotate query cells at multiple levels of resolution, and to visualize these cells in a 2D UMAP. After all datasets have been mapped, we subset all CD8 T cells (which encompass memory, naive, and effector states). We use monocle3 with default parameters^51^ to identify an integrated developmental trajectory that orders cells from all experiments.

#### Interpolating multiple histone modification profiles in single cells

In order to perform an integrated analysis of chromatin state at single-cell resolution, we first generate a set of single-cell profiles that comprise interpolated, genome-wide, and quantitative measurements of six histone modifications. In previous work^38^, we demonstrated how to utilize anchors between reference and query datasets to transfer not only discrete information (i.e. cell type annotations), but also quantitative measurements. For example, we demonstrated how to accurately interpolate cell surface protein measurements for a scRNA-seq dataset of 274,599 human bone marrow mononuclear cells generated by the Human Cell Atlas, based on a CITE-seq dataset of 35,543 cells from the same system.

We apply a similar workflow for this study (schematized in Figure 2A). First, we consider a CITE-seq dataset^30^, where each cell contains transcriptomic measurements alongside quantifications for a panel of 228 surface proteins. While the original dataset contains 161,764 cells, interpolating genome-wide profiles for multiple modalities at this scale creates memory and storage constraints, and we therefore select a random subset of 20,000 cells for downstream analysis. In Supplementary Figure 3B, we demonstrate that we obtain highly concordant results when repeating the procedure on a second random downsample of 20,000 cells.

We then find anchors between each scCUT&Tag-pro dataset, and the 20,000 CITE-seq cells. Anchors are determined using the FindTransferAnchors function, and leverage the same protein sPCA transformation as described above (‘Reference mapping’). We utilize these anchors to interpolate (i.e. ‘impute’) the number of fragments for each histone modification, in each 200bp genomic window, at single-cell resolution (see Stuart et al. 2019^38^, ‘Feature Imputation’). For example, in each of the 20,000 CITE-seq cells, the interpolated levels of H3K4me1 represent a weighted average of the anchor-forming cells in the H3K4me1 scCUT&Tag-pro. We repeat this procedure for each of the six modifications, as well as the ASAP-seq dataset^28^. Therefore, each of the 20,000 cells contains information for nine modalities, encompassing both the original measurements (RNA + protein), as well as seven interpolated modalities representing histone modifications and chromatin accessibility.

Based on our pseudobulk saturation analysis (Figure 1E), we used a weighted average of 500 anchors when interpolating values for each cell. We highlight that this weighting prioritizes anchors that are most similar to the query cell, enabling us to accurately interpolate histone modification profiles even for cell types present at moderate abundance (Figure 3C, Supplementary Figure 3A). However, for rare cell types, there is a possibility that an insufficient number of anchors may reduce the accuracy of these predictions. We therefore removed cell types that were present at <2.5% frequency in our reference dataset from further analysis.

In Figure 3B we plot the correlation between interpolated and original values for the H3K4me1 histone modification. Each point represents a 200bp genomic window. To avoid overplotting, we only plot windows on chr22 (n=254,093).

### Assigning chromatin states at single-cell resolution

In order to integrate information from multiple histone modifications we utilize the multivariate Hidden Markov Model introduced in ChromHMM^12^. For clarity, we reintroduce the key components of the model here. The model assumes that each 200bp genomic window (‘genomic interval’) can be represented based on one of *K* hidden states, where each state is defined by the combinatorial presence or absence of multiple histone modifications. The full likelihood of the model is:

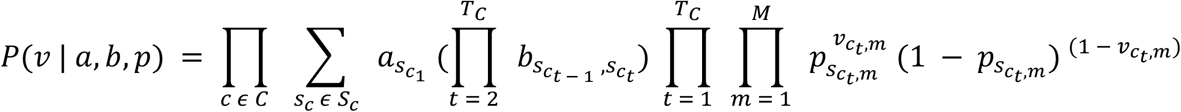

Where *v* represents the observed data, such that 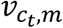 is ‘1’ if mark *m* is present at genomic interval *c*_*t*_ and ‘0’ otherwise. *p*_*k,m*,_ represents the probability that mark *m* is present at a genomic interval in state *k* (‘emission probability’), *b*_*i,j*_ represents the probability of the markov chain transitioning from state *i* to state *k* (‘transition probability’), and *s*_*c,t*_ represents the ‘hidden state’ associated with each at genomic interval *c*_*t*_.

The first step of the procedure is to learn a set of possible chromatin states, and their associated emission and transition probabilities. We use ChromHMM’s variant of the Baum Welch algorithm^12^ as implemented in the LearnModel command, setting the number of states to 12, and the maximum number of iterations to 1,000. As input to ChromHMM^17^, we provide the histone modification profiles for all cell types, where cell type identity was determined after reference mapping (level 2 resolution). The 12 learned states and associated emission probabilities are shown in Figure 3A.

After learning a set of chromatin states, we next infer the path of hidden states (hidden state sequence) through each chromosome, at single-cell resolution. To do so, we extend the ChromHMM method in order to calculate the posterior distribution of state assignments for each 200bp genomic interval in each cell. We refer to the following procedure as scChromHMM:

1. The input to scChromHMM is the set of interpolated histone modifications in single cells, as described above
2. As ChromHMM models multiple chromatin marks as an independent set of Bernoulli random variables, we must binarize the interpolated values present in each cell. As shown in Supplementary Figure 4A, the distribution of abundances for each histone modification exhibited clear bimodality. We assigned binarization thresholds via manual inspection (red dashed line), which were congruent with the results of Hartigan’s dip test. Binarization thresholds were set independently for each mark but kept constant across single cells. After thresholding, each genomic interval in each cell is represented by the presence or absence of six histone modifications.
3. We run the forward-backward algorithm on each cell independently, a dynamic programming algorithm to learn the posterior probabilities of hidden state variables in a hidden markov model^36^. The output of this procedure is a posterior probability distribution over all 12 states for each genomic interval in each cell.

### Comparison of scChromHMM and ChromHMM results

In addition to scChromHMM, we also annotated genomic intervals using the standard ChromHMM workflow^17^. While this approach does not return state predictions in individual cells, it does return posterior probability distributions for each cell type (level 2 resolution). We therefore compared the predictions from ChromHMM^17^ with the predictions from scChromHMM after grouping by celltype. In Figure 3E we perform a meta-analysis across all regions assigned to promoter, enhancer, repressor, and heterochromatic states, and explore the level of enrichment for each modification in a 10kb window centered on these regions. We combined the posterior probabilities for functionally related states (promoters: state 1 and 2; enhancers: state 3 and 4; repressor: state 9 and 10).

We compare results on CD14 monocytes, as this is the most abundant cell type in our dataset. For ChromHMM^17^, we selected all regions with >75% probability posterior probability in CD14 monocytes for meta-analysis, and for scChromHMM we selected all regions where the majority of CD14 monocytes had >75% posterior probability. In Supplementary Figure 4B, we also compare average PolII fragment abundance as measured by scCUT&Tag-pro on promoter regions as identified by either ChromHMM^17^ or scChromHMM.

### Identifying heterogeneity in chromatin state across a continuous trajectory

In previous work^38^, we used the Moran’s I statistic measure of spatial autocorrelation to identify genes whose expression varied as a function of spatial location. Here, we use a similar approach to identify 14,585 genomic intervals whose posterior probability (repressive state) varies across a continuous trajectory spanning Naive to Effector states in CD8 T cells (Figure 4A). We extract 3,157 CD8 T cells where we have calculated posterior state probabilities using scChromHMM, and where we also have obtained pseudotime estimates (see ‘Reference Mapping’). We use the pseudotime values to construct a Gaussian similarity kernel with unit bandwidth, which defines the spatial weights associated with the Moran’s I calculation. We calculate Moran’s I for each genomic interval as previously described^38^, and acknowledge the Trapnell Lab Monocle 3 tutorials for suggesting the use of Moran’s I to estimate spatial autocorrelation in single-cell data. We considered all intervals with Moran’s I > 0.15 as varying across the trajectory. In Figure 4A, we visualize the posterior probabilities of these regions as a function of pseudotime, after applying a moving average filter (k=21).

### Using chromVAR to identify associations between motifs and functional states

The chromVAR package^40^ identifies associations between transcription factors and chromatin accessibility by examining deviations (i.e. gains/losses) in the accessibility of peaks that share the same DNA sequence motif while carefully controlling for technical biases. Here, we aim to detect associations between transcription factors and the acquisition of distinct chromatin states. We therefore ran the chromVAR algorithm^40^, but instead of passing single-cell chromatin accessibility profiles as input, we utilized our single-cell posterior state probabilities. We repeat this process separately for promoter, enhancer, and repressor states. Using the promoter state as an example, chromVAR calculates - for each motif and each cell - the difference between the observed promoter posterior probabilities for peaks containing the motif, and the expected posterior probability (based on an average across all cells). chromVAR next performs a similar calculation for a matched background peak set in order to calculate bias-corrected deviation scores (Figure 4B, Supplementary Figure 4C).

### Characterizing regulatory priming based on repressive chromatin state

In Figure 4, we analyze cells based on their posterior probability (repressive state) at 19,306 transcriptional start sites (TSS). Based on a previous TSS definition^47^, we consider all genomic intervals that overlap with a window starting 2,500bp upstream of the TSS, and extending 500bp downstream. In each cell, we quantify the level of repression at each TSS as the average of the repressive state posterior probability for each of these windows. We generate a repressive score matrix that represents the level of repression at each cell (rows), at each TSS (columns). We apply TF-IDF normalization^50^ to this matrix followed by SVD, and use singular vectors 2:30 as input to UMAP visualization in Figure 4C.

In Figure 4D, we visualize cell-cell correlations based on the repressive score matrix. In the left heatmap, we use all 19,306 TSS as input to LSI, and use dimensions 2:30 to calculate a correlation matrix. In the right heatmap, we perform the same procedure, but exclude the top 3,000 TSS that were identified as highly variable based on gene expression levels, as determined by SCTransform^52^. In both heatmaps, cells are grouped based on their predicted annotation, but the order of cells within an individual group is random.

In Figure 4E, we consider the overlap between heterogeneity in repressive chromatin state and transcriptional variation. Using the FindMarkers command in Seurat (default parameters), which performs a wilcox sum rank test, we compared CD14 Monocytes to all other cells. We performed tests for scChromHMM-derived posterior probabilities (repressive state), H3K27me3 levels (as measured by scCUT&Tag-pro), and gene expression (as measured by CITE-seq), and kept all results with adjusted p-value < 0.01. To ensure that we robustly identified regions varying in their repressive state, we intersected the lists of TSS identified as differential by scChromHMM as well as H3K27me3, resulting in 1,597 loci. Of the associated genes, 257 were identified as differentially expressed in the transcriptome.

### Mapping scCUT&Tag datasets to a multimodal reference

In Seurat v4^30^, we utilize ‘supervised PCA’ (sPCA) to integrate multimodal (CITE-seq) and unimodal (scRNA-seq) datasets. The sPCA reduction aims to project transcriptomic measurements into a low-dimensional space that most closely resembles the WNN graph. We construct a weighted nearest neighbor graph that identifies, for each cell, a set of neighbor cells in the most similar molecular state based on a weighted combination of RNA and protein modalities. This graph represents a linear kernel, *L*, that can be used to supervise the dimensional reduction process for transcriptomic measurements.

As described in (Barshan et al, 2011)^46^, there is a closed form solution to sPCA based on a data matrix *X*, kernel *L*, and centering matrix *H*. The projection matrix *U* is represented by the eigenvectors of matrix *XHLHX*^*T*^. We apply the resulting projection matrix (*U*_*rna*_) to new scRNA-seq datasets, in order to identify anchors between query and reference datasets. We can repeat the same process using the protein modality, in order to calculate (*U*_*ADT*_).

Here, we apply a conceptually similar procedure to the integration of multimodal (H3K27me3 scCUT&Tag-pro, from this study) and unimodal datasets (H3K27me3 scCUT&Tag from Wu et.al.)^19^. For unsupervised analysis of the scCUT&Tag dataset, we downloaded fragment files from Gene Expression Omnibus with accession code GSE157910. We filtered cells with less than 250 fragments or more than 5,000 fragments, and repeated the LSI-based analysis workflow to perform unsupervised analysis (Figure 5A).

For each scCUT&Tag-pro dataset, the reference mapping procedure described above leverages the protein measurements to map each cell into a reference-defined space. We construct a k-nearest neighbor graph (*k=30)* in this space. This defines a kernel matrix *L*, where *L*_*i,j*_ *=* 1 if cells *i, j* are neighbors and 0 otherwise. We compute the matrix *XL*, where X is the original peak-barcode matrix that represents the quantified scCUT&Tag profiles in the multimodal experiment. We use the *XL* matrix as input to TF-IDF normalization and SVD. The result is a modified LSI that represents a low-dimensional transformation of the chromatin profiles, but has been supervised by the protein measurements. We use the FindTransferAnchors (reduction = ‘lsiproject’, dims = 1:30), to identify anchors between our scCUT&Tag-pro dataset and the unimodal query. We use these anchors to annotate cell types in the query dataset and assign prediction scores.

Lastly, we also use these anchors to interpolate protein values for 173 surface proteins in the scCUT&Tag query dataset. We then apply the previously computed *U*_*ADT*_ transformation to map these cells onto the original CITE-seq reference^30^. We perform this step for visualization purposes only, as it allows us to view the query scCUT&Tag dataset in the same UMAP visualization as Figure 2B, which represents all datasets in this study (Figure 2B).

