## Supplementary Figures for "Characterizing cellular heterogeneity in chromatin state with scCUT&Tag-pro"

### Supplementary Figure 1

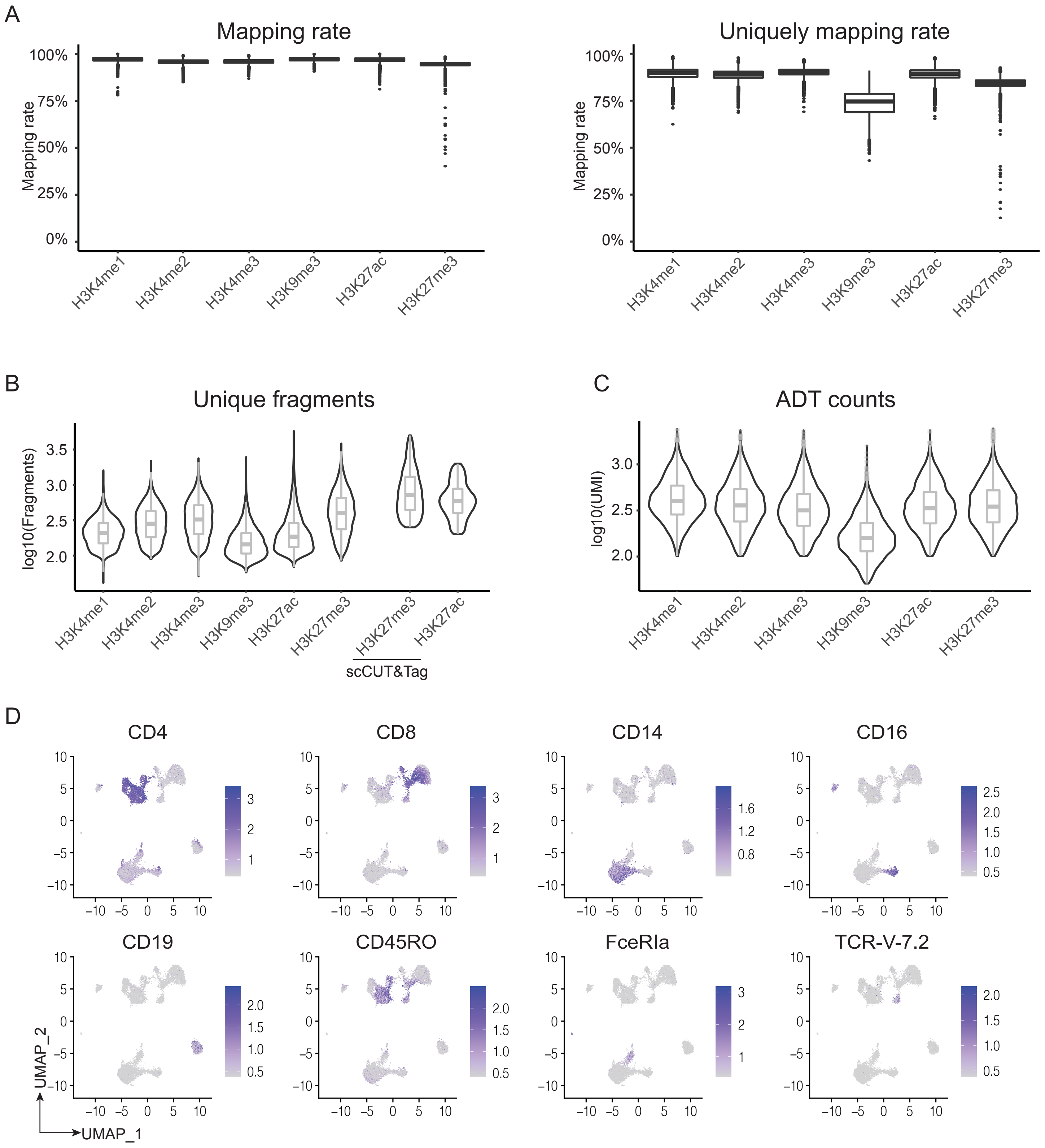


**Supplementary Figure 1: Quality control for scCUT&Tag-pro experiments.**

**(A)** Boxplot of single cell mapping rates and unique mapping rates for scCUT&Tag-pro experiments in PBMC. **(B)** Violin plots showing the number of unique fragments per cell detected in each PBMC scCUT&Tag-pro experiment, as well as from a recent publication describing scCUT&Tag in the same system (Wu et al, 2021). **(C)** Violin plots of unique antibody-derived tag (ADT) counts per cell for each experiment. **(D)** Feature plots visualizing the expression of canonical surface proteins for the H3K4me1 scCUT&Tag-pro experiment. Proteins visualized include markers of CD4 and CD8 T, CD14 and CD16 monocytes, B cells (CD19), Memory T (CD45RO), cDC2 (FceRIa), and MAIT (TCR-V-7.2) cells.

### Supplementary Figure 2

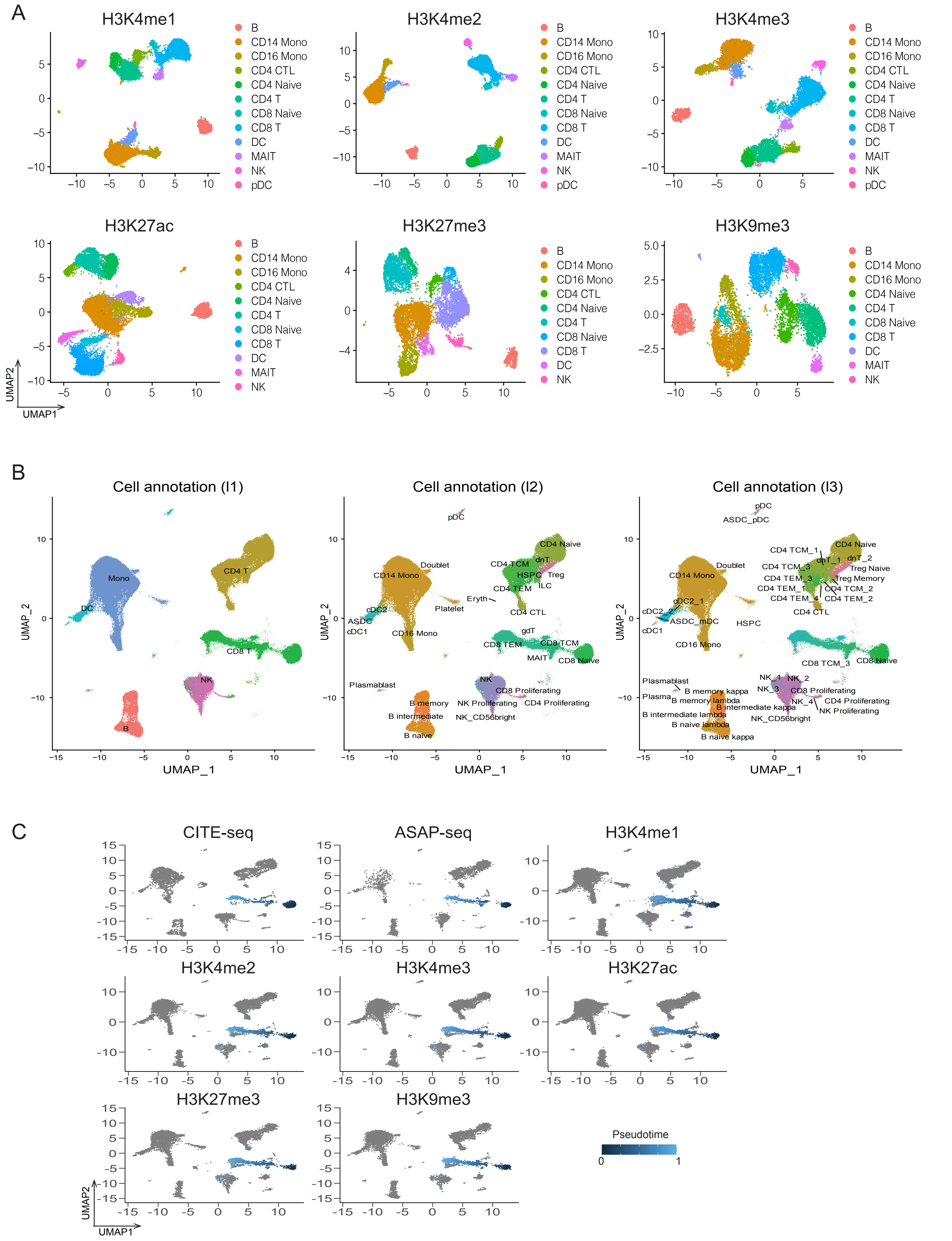


**Supplementary Figure 2: Multimodal and reference-based analysis of scCUT&Tag-pro**

**(A)** Weighted Nearest Neighbor (WNN) analysis of each of the six scCUT&Tag-pro datasets. Cells are visualized and clustered based on a weighted combination of the chromatin and cell surface protein measurements. **(B)** Cells from all experiments were mapped onto the reference CITE-seq dataset from (Hao et al, 2021). Shown are the reference-assigned labels for all cells. Same as Figure 2B, but visualizing all three levels of resolution. **(C)** Visualization of integrated developmental trajectory for CD8 T cells across all experiments. Each panel visualizes cells from a separate experiment, and CD8 T cells are colored based on their order along the integrated developmental trajectory.

### Supplementary Figure 3

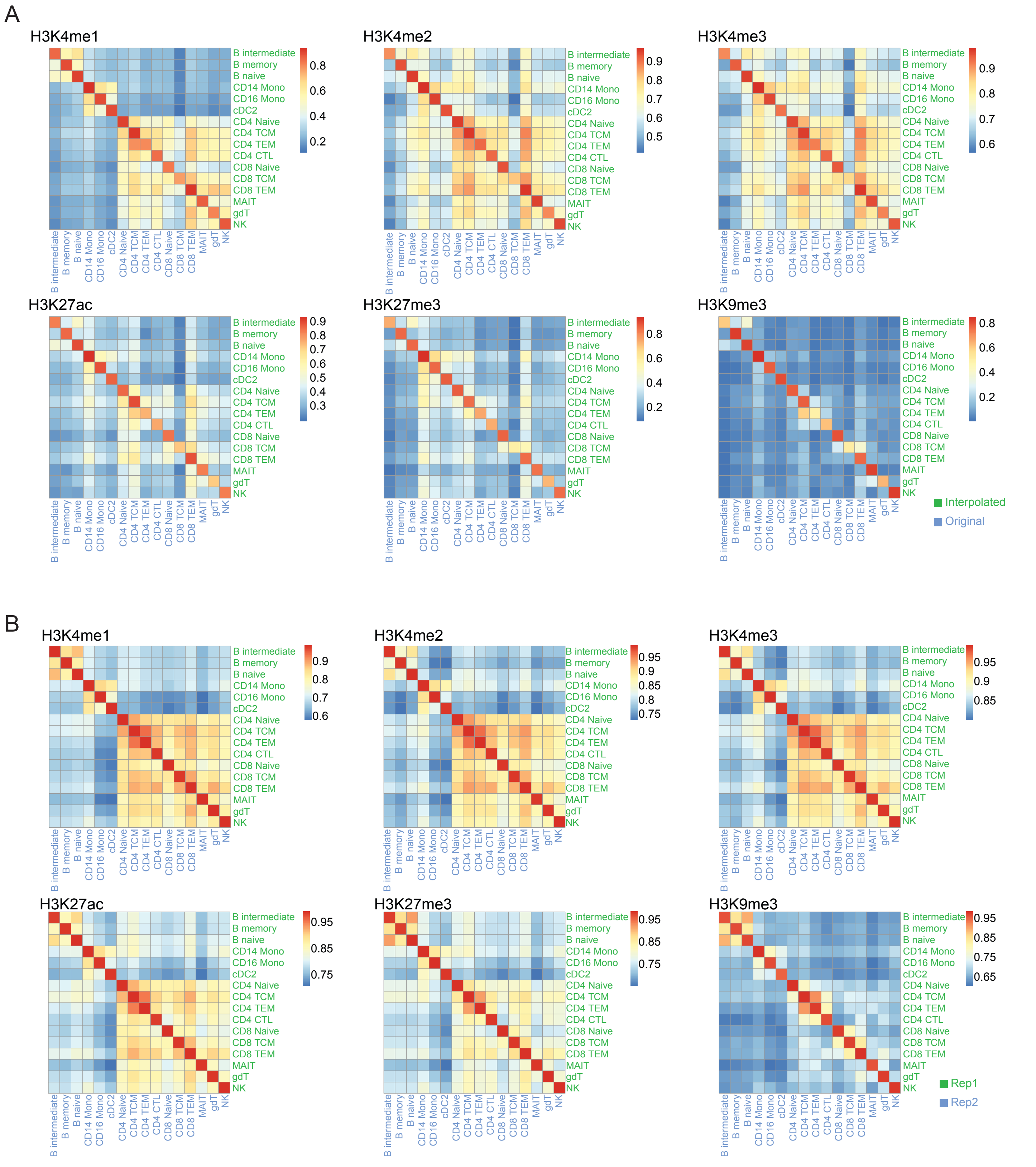


**Supplementary Figure 3: Accurate and robust interpolation of histone modification profiles.**

**(A)** Correlation matrix showing the relationship between original and interpolated values across all cell types. Same as Figure 3B, but for each of the histone modifications measured by scCUT&Tag-pro. **(B)** We repeated the interpolation twice, each time interpolating values into different random samples (n=20,000) of CITE-seq cells. Correlation matrix comparing the interpolated values from both replicates, demonstrating that we obtain highly concordant results.

### Supplementary Figure 4

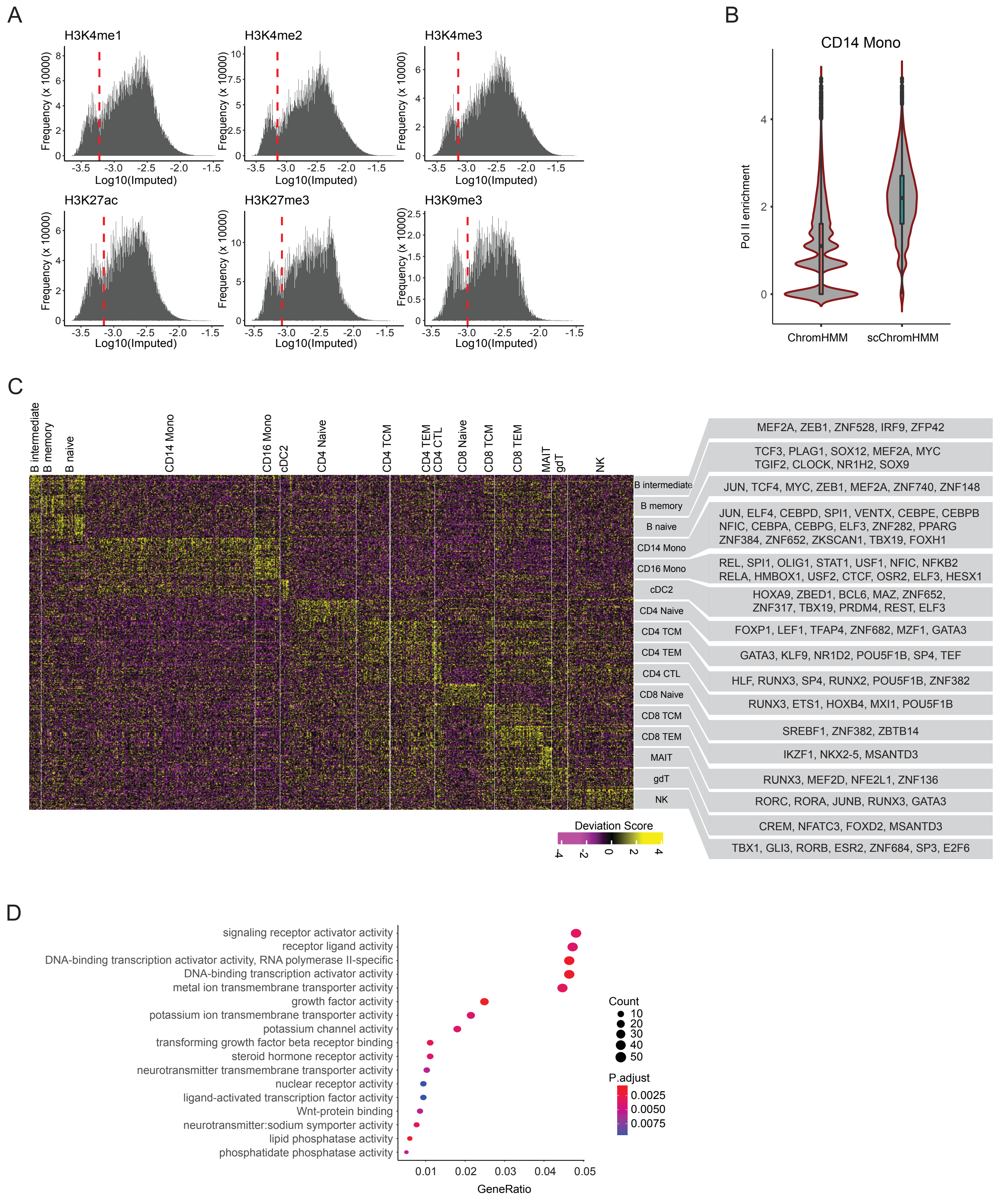


**Supplementary Figure 4: Inference and annotation of chromatin state at single-cell resolution**

**(A)** Histograms of interpolated values across single cells for each of the histone modifications measured by scCUT&Tag-pro. In each case, we observe a bimodal distribution. Red line marks the threshold used for data discretization, as required by ChromHMM and scChromHMM. **(B)** Violin plot showing the enrichment of Ser2/Ser5-phosphorylated RNA Polymerase II at active promoters identified by ChromHMM and scChromHMM in CD14 monocytes. **(C)** Heatmap showing heterogeneity in ChromVar deviation scores in single cells, grouped by their reference-derived annoatation. We used the scChromHMM-derived posterior probabilities (promoter state) as input to ChromVar, instead of chromatin accessibility levels. In grey boxes, we highlight each identified motif that is also differentially expressed at the transcriptional level. **(D)** Enriched GO terms for the 1,340 genes shown as blue points in Figure 4E.
